# High-Throughput Targeted Drug Screening for NF1-associated High-Grade Gliomas with ATRX Deficiency

**DOI:** 10.1101/2024.12.14.628525

**Authors:** Swati Dubey, Simran Rai, Fabiola Guillen, Ajeeth Iyer, Su Aung, Ming Yuan, Charles G. Eberhart, Fausto J Rodriguez

## Abstract

Neurofibromatosis type 1 (NF1)-associated high-grade gliomas (HGGs) harboring ATRX mutations exhibit an aggressive clinical phenotype, driven by heightened genomic instability and metabolic reprogramming. Existing therapies, including chemotherapy and radiotherapy, are limited by resistance mechanisms and formation of secondary malignancy, underscoring the need for novel therapeutic strategies. Here, we report the results of a high-throughput screening of 10,000 small molecules aimed at identifying compounds selectively targeting vulnerabilities associated with concurrent ATRX and NF1 loss. Among the screened compounds, K784-6195 (ChemDiv ID) emerged as a promising candidate, exhibiting marked selective cytotoxicity in NF1-associated glioma cell lines with ATRX deficiency (IC50 = 4.84 µM). In comparison, wild-type ATRX sporadic glioma cell lines (U251) exhibited significantly reduced sensitivity to K784-6195 (IC50 = 37.03 µM). However, ATRX knockout U251 glioma cells recapitulating concurrent ATRX and NF1 loss exhibited heightened susceptibility to K784-6195 (IC50 = 20-23 µM) compared to their wild-type counterpart. Metabolomic analysis revealed that K784-6195 treatment impairs metabolic pathways, including the pentose phosphate pathway, glutamine metabolism, and redox homeostasis, leading to oxidative stress and impaired cell survival. These findings highlight K784-6195 as a promising candidate for therapeutic development, offering a targeted approach for the treatment of NF-1 associated HGGs with ATRX deficiency.

## Introduction

Neurofibromatosis type 1 (NF1) is a genetic condition, conferring individuals to the predisposition of central and peripheral nerve tumorigenesis, wherein certain subsets exhibit a substantially aggressive phenotype often associated with ATRX mutations (Yuan et al., 2022; Romo et al., 2023; Dubey et al., 2024). The co-occurrence of ATRX and NF1 mutations is also detected in subsets of high-grade sporadic gliomas (Bedics et al., 2023). We have previously reported an association of ATRX loss with malignant gliomas in NF1 patients (Rodriguez et al., 2019) and have demonstrated therapeutic sensitivity to ATR inhibition in malignant gliomas with concurrent NF1 and ATRX deficiency, both in vitro and in vivo (Yuan et al., 2022).

Neurofibromin, the protein encoded by the NF1 gene, acts as a GTPase-activating protein (GAP) for RAS, converting active RAS-GTP into its inactive RAS-GDP form, thereby negatively regulating the RAS pathway (Tao et al., 2020). Since NF1 patients have nonfunctional neurofibromin, RAS is always active, which leads to constant activation of its downstream signaling pathways such as MAPK/ERK and PI3K/AKT/mTOR (Malone et al., 2014). The hyperactivation of these pathways promotes cell proliferation, metabolic reprogramming, immune evasion, and resistance to apoptosis, thereby contributing to tumor aggressiveness (Pungsrinont et al., 2021; Watterson & Coelho (2023); Cortes Ballen et al., 2024). Moreover, the role of neurofibromin in DNA repair underscores its loss as a driver of genomic instability, accelerating malignancy and the aggressive phenotype of NF1-associated tumors (Pemov et al., 2010; Mo et al., 2022). In the sporadic, or somatic, cases however, NF1 loss can be seen as an acquired mutation in different cancers (Philpott et al., 2017). This phenotype is often treatment resistant and more aggressive as well.

ATRX mutations, often co-occurring with NF1 loss, define a distinct molecular signature observed in clinically aggressive gliomas, including diffuse midline gliomas and pediatric high-grade gliomas (Lucas et al., 2022). ATRX deficiency is strongly linked to the alternative lengthening of telomeres (ALT) pathway, enabling telomere maintenance independent of telomerase (Amorim et al., 2016; Brosnan-Cashman et al., 2018; Pladevall-Morera et al., 2022). Activation of ALT, coupled with chromatin remodeling defects from ATRX loss, contributes to heightened genomic instability and an aggressive tumor phenotype. In NF1 associated tumors, this is exacerbated even further due to neurofibromin loss, compounding the aggressive phenotype (Rodriguez et al., 2016; D’Angelo et al., 2019).

The therapeutic landscape for NF1-associated high-grade gliomas remains challenging. The persistent activation of RAS/MAPK signaling confers resistance to standard chemotherapies (Tao et al., 2020). Mutations like TP53 and ATRX which often accompany NF1 loss can also lead to complications in chemotherapy as the tumor is better equipped to repair DNA damage caused by it. Additionally, radiotherapy poses significant risks for NF1 patients due to the potential development of secondary malignancies (Grill et al., 2009). These factors highlight the critical need for novel and targeted therapeutic approaches for treating NF1 associated high grade gliomas.

To address this unmet need, we conducted a high-throughput (HT) drug screening of 10,000 small molecules to identify compounds targeting the therapeutic vulnerabilities associated with concurrent NF1 and ATRX deficiencies in NF1-associated high-grade gliomas. Through comprehensive in vitro analyses, we prioritized compound K784-6195 (ChemDiv ID) based on marked selective efficacy, with a focus on advancing targeted therapies for this challenging clinical context.

## Materials and Methods

### Cell lines and culture conditions

U251, initially derived from an adult glioblastoma patient, was purchased from the American Type Culture Collection (ATCC). U251 cells were cultured in DMEM/F-12, GlutaMAX™ supplement (Gibco™ # 10565018) added with 10% FBS. U251 ATRX^−/−^ and U251 ATRX^−/2.02^, knockouts were utilized as previously generated in our study (Yuan et al., 2022). NF-1 associated high grade glioma cell line, JHH-NF1-GBM1, previously stablished in our lab (Yuan et al., 2022) was grown in DMEM/F12 containing 2mM GlutaMax (Gibco™ #35050061), 1X B27 supplement (Gibco™ #12587010), 20 ng/mL EGF (PeproTech, Cranbury, NJ, USA), 20 ng/mL FGF-b (PeproTech), 5 µg/mL Heparin (Millipore SIGMA, Burlington, MA, USA). All cells were grown in a 37°C humidified incubator with 5% CO_2_. Regular mycoplasma testing was performed, and human cell line identities were confirmed by short tandem repeat (STR) profiling.

### High-throughput drug screening

High-throughput drug screening was conducted using an automated liquid handling system to ensure precise and consistent cell dispensing across a 384-well plate format. U251 and its ATRX knockout (KO) cells were seeded at 100 cells per well, while JHH-NF1-GBM1 cells were seeded at 500 cells per well in 50 μL of cell culture media. The plates were incubated overnight in a humidified incubator at 37°C with 5% CO_2_ to facilitate proper cell attachment. The following day, a library of 10,000 diverse compounds received from a well-established NIH Molecular Libraries Probe Centers Network (MLPCN), Johns Hopkins ChemCORE was screened, with each compound administered at a final concentration of 1 μM by dispensing 5 nL of a 10 mM compound stock solution into each well using an automated liquid handler optimized for high-throughput dispensing. Plates were incubated for 5-7 days (5 days for U251 and ATRX KO cells, 7 days for JHH-NF1-GBM1 cells) to allow the compounds to exert their effects. To assess the impact of the compounds on cell viability, a CellTiter-Blue Cell Viability Assay (Promega) was performed according to the manufacturer’s protocol and fluorescence was measured at excitation/emission wavelengths of 560 nm/590 nm using a BMG CLARIOstar multimode microplate reader. Cell viability for each well was calculated using the following formula:

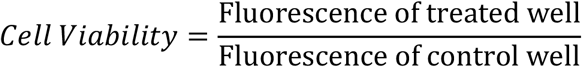

Control well represents the cells treated with vehicle (DMSO). RStudio was used for further data processing and visualization.

### Cell viability assay and IC50 determination

To evaluate cell viability and determine the IC50 values of selected compound(s), a PrestoBlue Cell Viability assay was performed. Optimal seeding densities were established as 1,000 cells per well for U251 cells and 10,000 cells per well for JHH-NF1-GBM1 in 96-well plates. Cells were manually dispensed into each well in 100 µL of culture media. Plates were incubated at 37°C in a humidified atmosphere with 5% CO_2_ to allow cell attachment. Candidate compound(s) were prepared in serial dilutions and added to the wells in 100 µL of culture media, resulting in a total well volume of 200 µL/well. Plates were incubated for five days under the same conditions. Following incubation, 20 µL of PrestoBlue reagent was added to each well, and plates were protected from direct light during the reaction. After 2 hours, fluorescence was recorded using Molecular Devices SpectraMax M5 microplate reader at excitation/emission wavelengths of 560 nm/590. Fluorescence data were normalized to DMSO-treated controls, and cell viability was calculated. Nonlinear regression analysis was conducted using GraphPad Prism to generate dose-response curves and determine IC50 values for each compound. Comparisons of IC50 values were made across cell lines to assess compound efficacy and selectivity.

### Metabolite labeling and extraction for glucose and glutamine tracing

Cells were seeded in six-well plates at a density of 700,000 cells per well in routine culture medium and incubated overnight at 37°C with 5% CO_2_. For glucose and glutamine-labeling experiments, SILAC Advanced DMEM/F-12 Flex Media (Gibco™ #A2494301) devoid of glucose and glutamine, supplemented with 17.5 mM ^13^C-glucose (Sigma Aldrich #660663), 4.5 mM ^15^N-glutamine (MedChem Express #HY-N0390S), 0.69 mM L-arginine (Thermo Scientific Chemicals #A15738.14), 0.49 mM L-lysine (Thermo Scientific Chemicals #J62225.14), 15 mM HEPES (Gibco™ #15630130), 1X B27 supplement (Gibco™ #A1486701), 20 ng/mL EGF (Peprotech #AF-100-15), 20 ng/mL FGF-b (Peprotech #100-18B), and 5 µg/mL heparin (Thermo Scientific Chemicals #A16198.03), was prepared. The specified drug treatment was added directly to this medium. The pre-existing culture medium was completely replaced with the prepared labeling medium, and cells were incubated for 24 hours to achieve steady-state labeling. Metabolites were harvested by transferring cells into polypropylene centrifuge tubes and centrifuging at 500–800 × g for 5 minutes to pellet the cells. A swing-bucket centrifuge was used to ensure gentle pelleting. After centrifugation, tubes were placed on ice, and the medium was carefully aspirated. The cell pellets were washed briefly by gentle resuspension in ice-cold PBS, followed by re-pelleting at 4°C. The wash solution was removed thoroughly without disturbing the pellet. Cell pellets were extracted with 1 mL ice-cold 80% methanol/20% water. The samples were vortexed for 30 seconds and incubated at −80°C for 30 minutes to facilitate metabolite extraction and protein precipitation. Following incubation, the samples were allowed to warm on ice, vortexed again, and centrifuged at 16,000 × g for 10 minutes at 4°C. The supernatants were transferred to glass vials and dried using an evaporator without heat (Genevac EZ-2 Elite, aqueous program). Dried metabolite extracts were stored at −80°C until further analysis by liquid chromatography–mass spectrometry (LC-MS).

### Metabolomics Analysis

Mass spectrometry-based metabolomics was conducted at the UCLA Metabolomics Core Facility. Dried metabolites were resuspended in 100 µL 50% ACN:water and 5 µL was loaded onto a Luna 3µm NH2 100A (150 × 2.0 mm) column (Phenomenex). The chromatographic separation was performed on a Vanquish Flex (Thermo Scientific) with mobile phases A (5 mM NH4AcO pH 9.9) and B (ACN) and a flow rate of 200 μL/min. A linear gradient from 15% A to 95% A over 18 min was followed by 7 min isocratic flow at 95% A and re-equilibration to 15% A. Metabolites were detected with a Thermo Scientific Q Exactive mass spectrometer run with polarity switching in full scan mode with an m/z range of 70-975 and 140.000 resolution. Maven (v 8.1.27.11) was utilized to quantify the targeted metabolites by area under the curve (AreaTop) using accurate mass measurements (< 5 ppm) and expected retention time previously verified with standards. C13 and N15 natural abundance corrections were made using AccuCor2. Data analysis was performed using in-house R scripts.

## Results and Discussion

### Down-selection of compounds targeting vulnerabilities from concurrent NF1 and ATRX loss

High-throughput drug screening was performed using a diverse library of 10,000 compounds on an NF1-associated ATRX mutant glioblastoma cell line (JHH-NF1-GBM1), U251 high-grade glioma cells with ATRX knockout (U251 ATRX KO), effectively mimicking the concurrent loss of ATRX and NF1, along with wild type U251 cells. Supplementary table 1 include percent survival for all three cell lines (JHH-NF1-GBM1, U251 wild-type (WT), and U251 ATRX KO) across the compound library. A diverse spectrum of drug responses was observed among the cell lines (Figure 1B). Specifically, compounds eliciting darker hues in the U251 WT column (indicative of higher survival rates) and lighter hues in the U251 ATRX KO and JHH-NF1-GBM1 columns (indicative of lower survival rates) identified 105 candidate compounds with potential selective activity against vulnerabilities associated with concurrent ATRX and NF1 loss. Selection criteria included a survival threshold of ≤30% in both lines, with minimal impact on the viability of U251 WT cells (Figure 1B). To further refine the candidate pool, we leveraged biological activity data from PubChem and target-binding interactions from BindingDB (Gilson and Tiqing (2023)). This integrative approach ensured the selected compounds were not only effective in survival assays but also demonstrated potential relevance to cancer-associated pathways and targets. This process ultimately narrowed the candidate list to eight compounds (ChemDiv IDs: [8014-9607], [D216-0413], [6521-0021], [8018-8707], [4817-0512], [D216-0413], [K784-6195], [D257-0773]), which were further validated at a high concentration (100 µM) for their ability to inhibit cell proliferation, particularly in the context of ATRX and NF1 loss (Figure 1C). To our surprise, only one compound, K784-6195, demonstrated significant inhibitory activity at a considerable concentration, making it the focus of our subsequent studies. The calculated IC50 values for K784-6195 revealed its selective potency in the presence of ATRX and NF1 deficiency.

**Figure 1.**
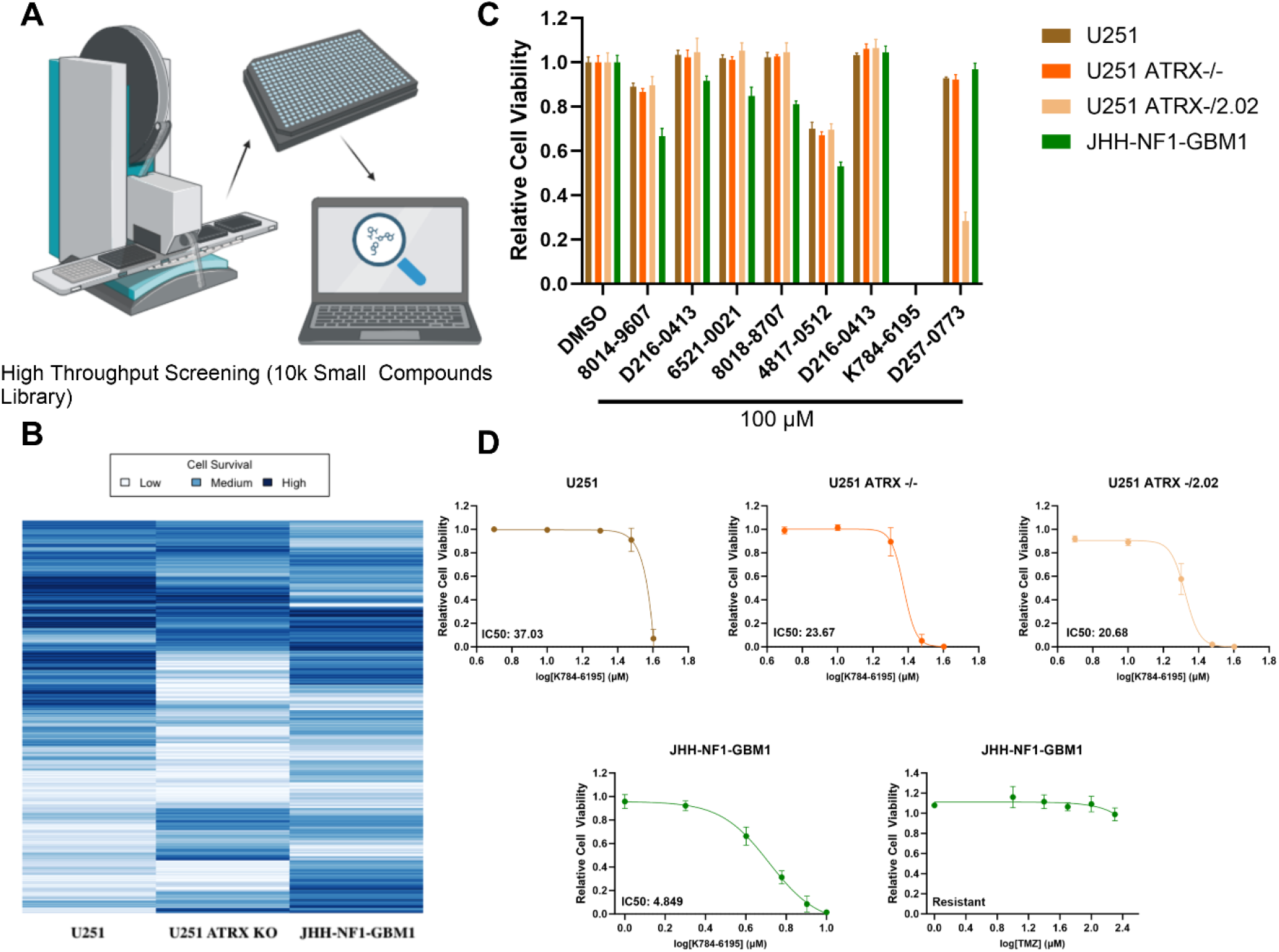
(A) High-throughput screening (HTS) of a 10,000-compound library. (B) Heatmap illustrating survival rates across U251 WT, U251 ATRX KO (-/-), and JHH-NF1-GBM1 cell lines, with a blue gradient indicating survival rates (dark blue: higher survival, light blue: lower survival). (C) Cell viability assessment of down-selected compounds, where cells were highly susceptible to K784-6195 at 100 µM concentration, while (D) at lower concentration range K784-6195, demonstrated marked susceptibility against NF1-associated HGG line. n=5, Mean±SD from three independent experiments.

Specifically, the NF1-associated high-grade glioma (HGG) cell line JHH-NF1-GBM1 exhibited the highest susceptibility to K784-6195, with an IC50 value of 4.84 µM. In comparison, the U251 WT and U251 ATRX KO cell lines showed IC50 values of 37.03 µM and 20-23 µM, respectively (Figure 1D). Notably, this compound demonstrated marked susceptibility against NF1-associated HGG line that was otherwise resistant to temozolomide (TMZ). This finding establishes a strong foundation for further investigations into the mechanisms of action of K784-6195, with the aim of developing more effective treatment options for NF1-associated gliomas, particularly in cases where existing therapies have limited efficacy.

### Metabolomic Insights into K784-6195 Activity in NF1-Associated HGGs

To elucidate the mechanistic basis of K784-6195 susceptibility in NF1-associated high-grade glioma (HGG) cells, we performed targeted steady-state metabolomics using isotopically labeled glucose and glutamine 24 hours post-treatment with K784-6195 at its IC50 concentration. The treatment led to significant alterations in metabolic pathways, as evidenced by isotopic tracing studies (Figure 2A, B) and global metabolomic profiling (Figure 2C).

**Figure 2.**
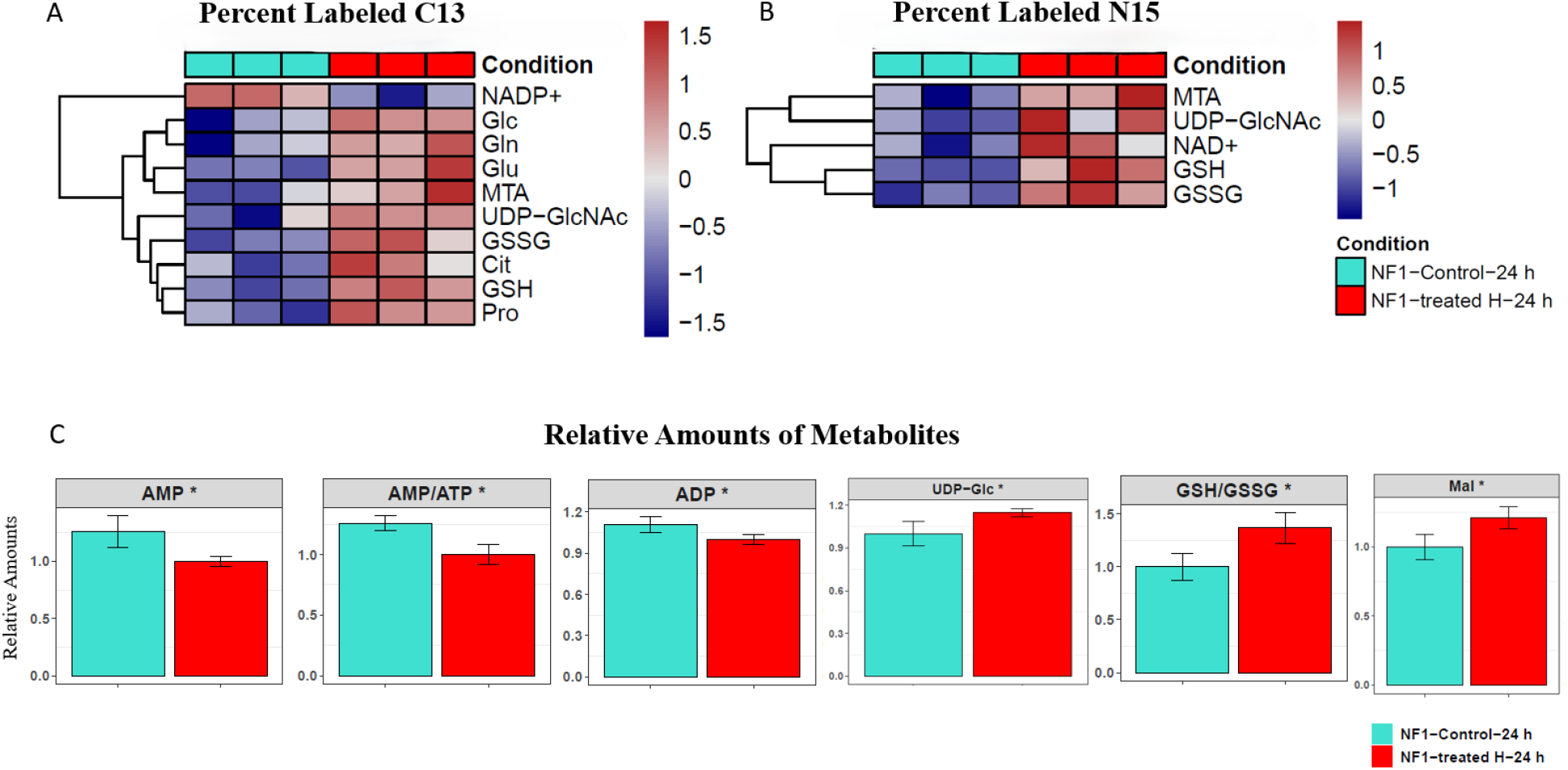
Steady-state metabolomic profiling of JHH-NF1-GBM1 cells after 24-hour treatment with K784-6195. Heatmaps show differentially regulated metabolites from (A) 13C-Glucose and (B) 15N-Glutamine isotopic labeling. (C) Global metabolite changes highlight treatment-induced metabolic alterations.

Post-treatment, the reduction in C13-labeled NADP^+^ levels (Figure 2A), derived from [C13]-labeled glucose, suggests impaired pentose phosphate pathway (PPP) activity or enhanced NADPH consumption in redox balance and anabolic biosynthesis. Concurrently, elevated oxidized glutathione (GSSG) alongside reduced glutathione (GSH) levels reflects oxidative stress induced by K784-6195 treatment. Elevated proline levels suggest its role as a stress response metabolite, stabilizing proteins and membranes under metabolic or oxidative stress (Liang et al., 2013). Increased labeling of UDP-N-acetylglucosamine (UDP-GlcNAc) points to upregulated hexosamine biosynthesis, a crucial adaptive response in stressed cancer cells, facilitating glycosylation of proteins and lipids to support signaling, proliferation, and survival (Chiaradonna et al., 2018).

Similarly, in parallel, enhanced incorporation of N15 from glutamine was observed in UDP-GlcNAc, GSSG, GSH, suggesting active glutamine-derived antioxidant defenses, balancing the elevated oxidative stress triggered by treatment. Further, elevated N15 labeling in NAD^+^ reflects increased reliance on glutamine for NAD^+^ biosynthesis, likely through the salvage pathway, which supports redox homeostasis and DNA repair under stress.

Despite these metabolic adaptations, cell viability was significantly reduced following treatment with K784-6195. This indicates that the concurrent loss of NF1 and ATRX sensitizes glioblastoma cells to perturbations in redox homeostasis. NF1 loss enhances RAS signaling, driving metabolic demand (Masgras & Rasola (2021), while ATRX mutations impair chromatin structure and DNA repair. Together, these vulnerabilities may exacerbate metabolic and oxidative stress under drug treatment, tipping the balance toward cell death.

Further, analysis of global metabolite levels (Figure 2C) showed significant upregulation of UDP-glucose in treated cells, pointing to metabolic shifts in the glucuronic acid pathway and its downstream extensions. Elevated UDP-glucose levels suggest potential inhibition of UDP-glucose 6-dehydrogenase (UGDH), the enzyme responsible for converting UDP-glucose to UDP-glucuronic acid. Previous studies in glioblastoma and lung cancer models have linked UGDH inhibition to reduced cancer cell proliferation and migration via glycosaminoglycan biosynthesis and stabilization of metastasis-promoting SNAI1 mRNA (Oyinlade et al., 2018; Wang et al., 2019). Further studies are underway to validate these findings and exploring the therapeutic potential of combining K784-6195 with other agents targeting redox and metabolic vulnerabilities in NF1-associated high-grade gliomas.

## Conclusions

Our high-throughput drug screening approach identified K784-6195 as a promising candidate compound selectively targeting vulnerabilities associated with concurrent NF1 and ATRX deficiencies in high-grade gliomas. The compound demonstrated significant inhibitory activity in NF1-associated glioma cell line, which was otherwise resistant to conventional treatments like temozolomide. Mechanistic insights from metabolomic profiling revealed significant alterations in redox homeostasis, underscoring the metabolic stress induced by K784-6195 treatment. Future work will explore the compound’s mechanism of action in greater detail and evaluate its efficacy in combination therapies to enhance therapeutic outcomes in NF1-associated gliomas.

## Acknowledgements

We thank the ChemCore High Throughput Facility at the Johns Hopkins School of Medicine for providing the ChemDiv Diverse Set 10k library used in our screening experiments. We acknowledge the UCLA Metabolomics Center for metabolomics analysis and extend our heartfelt gratitude to Dr. Johanna ten Hoeve-Scott for her invaluable assistance in planning the metabolomics experiments. We also thank the Vossel Lab at UCLA for granting access to their SpectraMax M5 microplate reader. Figure 1A was created using BioRender.com.

## Funding

This research was funded by the UCLA SPORE in Brain Cancer (P50CA211015).

## Notes

https://pubchem.ncbi.nlm.nih.gov/substance/505366702

### Competing Interest Statement

The authors have declared no competing interest.

### Summary of Updates

Funding Info updated; This research was funded by the UCLA SPORE in Brain Cancer (P50CA211015).

